# Modulation of intestinal bile acids influences colonic mucosal responses

**DOI:** 10.1101/2025.02.21.639473

**Authors:** Esther Wortmann, Tanja Groll, Anne Strigli, Kenneth Peuker, Colin Volet, Leona Arps, Rizlan Bernier-Latmani, Sebastian Zeissig, Katja Steiger, Moritz Middelhoff, Thomas Clavel

**Affiliations:** Functional Microbiome Research Group, Institute of Medical Microbiology, University Hospital of RWTH Aachen, Germany; Institute of Pathology, School of Medicine and Health, Technical University of Munich, Germany; Comparative Experimental Pathology, School of Medicine and Health, Technical University of Munich, Germany; Department of Internal Medicine A, University Medicine Greifswald, Greifswald, Germany; Environmental Microbiology Laboratory, School of Architecture, Civil and Environmental Engineering, École Polytechnique Fédérale de Lausanne, Lausanne, Switzerland; Center for Regenerative Therapies Dresden (CRTD), Technische Universität (TU) Dresden, Dresden, Germany

**Keywords:** gut microbiome, secondary bile acids, gut epithelial proliferation, colorectal cancer

## Abstract

**Background:** Elevated levels of secondary bile acids produced by the gut microbiome, in particular deoxycholic acid (DCA), influence epithelial cell proliferation and accelerate the development of colorectal cancer (CRC) under adverse dietary conditions, such as long-term, high fat intake. However, their effects on the intestinal epithelium have not been studied in detail.

**Aim:** To determine gut epithelial responses to bile acid modulation *in vivo* and *in situ*.

**Methods:** We performed targeted colonization of gnotobiotic mice followed by single-cell RNA sequencing (scRNA-Seq) of colonic epithelial cells combined with immunostaining of human biopsies from: (i) an observational patient cohort with hyperproliferative polyps or cancer; (ii) an interventional study with bile acid-scavenging drugs.

**Results:** Colonization of mice with a synthetic bacterial community together with the 7α-dehydroxylating species *Extibacter muris* resulted in DCA production. ScRNA-Seq of colonic epithelial cells revealed increased cell density of bile acid-sensitive enterocytes but fewer stem cells, goblet cells, and transit amplifying cells in mice exposed to DCA. This was associated with increased expression of pyruvate dehydrogenase kinase (*Pdk4*) and decreased expression of mucin (*Muc2*). PDK expression was also increased in human hyperplastic polyps and adenomas, whilst MUC2 expression was reduced in adenomas and carcinomas compared to normal mucosa. In addition, human exposure to bile acid sequestrants was associated with enhanced epithelial proliferation.

**Conclusion:** This study provides insight into intestinal epithelial cell responses to bile acids and their potential clinical relevance.

## Introduction

The human gut is colonized by complex communities of microbes known as the microbiota (1). Within the microbiota, gut bacteria play a causal role in a wide range of non-communicable diseases, including colorectal cancer (CRC) (2–6). One way in which gut microbes influence pathophysiological responses in the host is through the numerous metabolites they produce (7, 8). Among these metabolites, bile acids, secreted in conjugated form in the proximal small intestine, represent a broad spectrum of molecules at the host-microbe interface (9).

Primary bile acids are produced in the liver. However, they are further metabolized by bacteria in the intestine, which produce so-called secondary bile acids, for example by deconjugation and 7α-dehydroxylation (10). Microbial metabolism modifies the bioavailability and bioactivity of bile acids. In particular, 7α-dehydroxylation leads to the production of dominant secondary bile acids, including deoxycholic acid (DCA) and lithocholic acid, which are more hydrophobic, bind with a different affinity to bile acid receptors, and have been implicated in CRC. An important membrane-bound bile acid receptor is the G protein-coupled bile acid receptor 1 (GPBAR1, also referred to as TGR5). A recent study showed that bile acid binding to TGR5 in intestinal stem cells promotes regeneration of the intestinal epithelium after injury (11). In addition, TGR5 can activate mitogen-activated protein kinase (MAPK) signaling through several pathways such as extracellular signal-regulated kinases (ERK1/2), steroid receptor coactivator (SRC), and the mechanistic target of rapamycin (mTor) (12). Farnesoid X receptor (FXR)-dependent bile acid signaling has also been shown to regulate colonic epithelial wound healing and stem cell proliferation (13, 14).

Diet-microbiota interactions have been implicated in the development of CRC (15–17). Increased dietary fat intake over time is an important risk factor, and increased production of secondary bile acids by gut bacteria plays a role in this process (17–19). Although intestinal epithelium proliferation is closely associated with CRC development, the effects of secondary bile acids on epithelial cell responses have not been studied in detail *in vivo*. In this work we used gnotobiotic mice and single-cell RNA sequencing (scRNA-Seq) in combination with immunostaining of human colonic tissue to investigate epithelial responses to bile acid regulation.

## Methods

### Bacterial stocks for mouse colonisation

Fresh overnight cultures of each bacterial strain within the Oligo-Mouse Microbiota (OMM)12 (20) and of the DCA-producing species *Extibacter muris* DSM 28560^T^ (21, 22) were generated in an anaerobic rich medium (23), which contained: 18.5 g/L Brain Heart Infusion (Oxoid CM1135); 15 g/L Tryptone (Oxoid CM0129), 5 g/L yeast extract (Roth 2363.1); 2.5 g/L K2HPO4 (Supelco 105104); 1 mg/L hemin (Sigma-Aldrich H9039); 0.5 g/L glucose (Sigma-Aldrich G8270); 0.4 g/L Na2CO3 (Merck 106392); 1 mg/L resazurin (Fisher Scientific 10371053); 250 mg/L mucin (Sigma-Aldrich M2378); 0.5 g/L cysteine-HCl (Sigma-Adrich C7352); 0.5 mg/L menadione (Sigma-Aldrich M5625); 30 ml/L fetal calf serum (Sigma-Aldrich F7524, complement inactivated). The strains were mixed according to their OD_600_ value in sterile PBS supplemented with 10% glycerol and stored in serum bottles (Sigma-Aldrich, cat. no. Z113948) containing an anoxic gas mixture and sealed with a crimp cap (TH.GEYER, cat. no. 7608142). All strains were checked for purity using microscopy and Sanger sequencing of 16S rRNA genes. In addition, the culture mix was checked by 16S rRNA gene amplicon sequencing prior to inoculating the mice.

### Mice

The animal experiment was performed under LANUV ethical approval nr. 81-02.04.2018.A425 in accordance with EU regulation 2010/62/EU. Germfree *Apc*^1638N/+^ mice (B6/J.129-(Apc1638N)tm) were colonized with either, the synthetic microbial community OMM12 or OMM12+E (*i.e.*, OMM12 with the addition of *E. muris*). The germfree mice were transferred to sterile isocages (IsoCage P system, Tecniplast, Italy) and inoculated orally (50 μL) and rectally (100 μL) with the cryopreserved bacterial mixtures at the age of 5 weeks, with a second inoculation 2 days after the first gavage to favor the engraftment of strictly anaerobic species. The amount of gavage solution was adjusted to a maximum of 10% (v/w) body weight for mice weighing <15 g. After a stabilization period of 2 weeks, the standard chow feed (autoclaved, 120°C for 20 min) was changed to an experimental control diet (ssniff Spezialdiäten GmbH, cat. nr. S5745-E902) sterilized by irradiation with 2×25 kGy. To take sex and cage effects into account, one female and one male mouse from each colonization group (OMM12 and OMM12+E) were used for scRNA-Seq.

### Epithelial cell isolation

The entire colon of the two mice per colonization group was pooled and washed 3 times with ice-cold PBS. The tissue was cut into small pieces (2-5 mm) and transferred into 30 mL ice cold RPMI + 5% FCS (gibco, cat. nr. 31870-025; Sigma-Aldrich, cat. nr. F7524) containing 2 mM EDTA. The tubes were gently inverted by hand and incubated on a horizontal shaker (Universal Shaker SM 30 Edmund Bühler GmBH) at 110 rpm and room temperature for 15 min. The suspension was passed through a 100 μm cell strainer (Corning, cat. nr. CLS431752) and the filtrate was centrifuged (300 g, 5 min, 4 °C) and re-suspended in 500 μL PBS-FCS (5 %)-EDTA (0.01 mM) using wide bore pipette tips (VWR Europe, cat. nr. 613-0752) to collect the first fraction of isolated epithelial cells. This procedure was repeated and fractions 6-8 were then pooled, centrifuged (300 g, 5 min, 4 °C), re-suspended in 250 μL PBS-FCS (5 %)-EDTA (0.01 mM), and transferred to a 96-well plate (Greiner BioOne, cat. nr. 651201). After blocking with 10% normal rat serum (Dianova, cat. nr. 012-000-001), cells were stained with Epcam-APC (1:200, eBioscience™ cat. nr. 17-5791-82) and CD45-BV510 (1:200, BioLegend, cat. nr. 103138) for 20 min. To stain for dead cells, DAPI (1:200, Roth, cat. nr. 6335.1) was added just before the last centrifugation step (300 g, 5 min, 4 °C). The stained cells were filtered and sorted for single, live, CD45-negative, Epcam-positive cells with a BD FACSAriaTM Ilu, equipped with a 100 μm nozzle and kept at 4 °C in 5 mL PBS-BSA (1 %) (Merck, art. nr. SRE0036). The collected cells were counted and evaluated for quality and shape by staining with trypan blue and re-suspended to achieve a density of approximately 1,000 cells/µL.

### Single cell sequencing

Library preparation and quality control were done according to the Chromium Next GEM Single Cell 3’ Reagent Kits v3.1 protocol, CG000204 Rev D (10x Genomics, Inc., United States), aiming for an output of 10,000 cells and a sequencing depth of 20,000 read pairs per cell. RNA sequencing was performed on an Illumina NextSeq 550 (paired-end, 2 × 75 bp) at the IZKF Core Facility Genomics, University Hospital of RWTH Aachen. For the OMM12 sample, an estimated number of 8,161 cells were sequenced, with 32,940 mean reads per cell, 4,537 median UMI counts per cell, 1,121 median genes per cell, and a sequencing saturation was 36.9%. For the OMM12+E sample, an estimated number of 5,950 cells were sequenced, with 51,998 mean reads per cell, 7,144 median UMI counts per cell, 1,544 median genes per cell, and a sequencing saturation of 41.8 %.

### Singe-cell sequencing analysis

Reads were aligned to the mouse genome (mm10-3.0.0) using the Cell Ranger pipeline (version 6.1.1). Sample integration, creation of a Seurat object, and filtering were done using a scRNA pipeline (https://github.com/CostaLab/scrna_seurat_pipeline). Data was log-normalized with a scale factor of 10,000 and centered with a scale maximum of 10. Cells with a number of genes (nFeature_RNA) <400 and number of molecules per cell (nCount_RNA) >40,000 were filtered out. Effects of cell cycling (combined effects of the calculated G2M.Score and S.Score), mitochondrial (percent.mt) and ribosomal genes (percent.ribo) were regressed out for cell clustering. Genes from the X-inactivation center (Xist, Tsix, Jpx, Ftx) were filtered out before clustering to reduce sex-specific effects. Unsupervised clustering was performed using a shared nearest neighbors graph with a resolution of 0.3. Downstream analysis was done with the Seurat R package version 4.1.1 [4–7] using “RNA” as the default assay. Visualization of the clusters as UMAPs was done with the function “DimPlot” using reduction = “INTE_UMAP”. Cell cluster identification was done using cell type markers proposed previously (24). Proportional differences between the colonization groups was done by using a permutation test provided by the R package ‘scProportionTest’ (version v1.0.0) available at https://github.com/rpolicastro/scProportionTest (25). Cell clusters were analyzed separately for differential gene expression analysis with the Seurat function Findmarkers using the Wilcoxon test and default settings. The highest number of molecules and genes were detected in cell clusters 5, 7, and 8, making them the most transcriptionally active cell populations in this dataset (**Supplementary Figure S1**). The percentage of mitochondrial genes, which can be an indicator of dying cells, was 25 % and 26% in OMM12 and OMM12+E mice, respectively, driven mainly by cell cluster 3 and 6 (**Supplementary Figure S1**). Ribosomal gene expression, considered as an indicator of cell proliferation (26), was cell type specific, with an average of 6.3 % and 3.4 % ribosomal genes detected in OMM12 and OMM12+E mice, respectively.

### 16S rRNA gene amplicon analysis

Metagenomic DNA isolation from colon content and 16S rRNA amplicon sequencing were performed as described before (27), resulting in an average of 11,718 ± 3,639 reads per sample. Sequences were identified by alignment to the reference 16S rRNA gene sequences of OMM strains and *E. muris* or by using EzBioCloud (28).

### Bile acid analysis

Bile acids were quantified in cecal content from the gnotobiotic mice stored at −80 °C as described previously (17). Briefly, the samples were extracted by solid phase extraction and measured using an Agilent ultrahigh-performance liquid chromatography 1290 series coupled in tandem to an Agilent 6530 Accurate-Mass Q-TOF mass spectrometer 66 equipped with a Zorbax Eclipse Plus C18 column (2.1 × 100 mm, 1.8 μm) and a guard column Zorbax Eclipse Plus C18 (2.1 × 5 mm, 1.8 μm). All parameters and running conditions were as before (17). In total, 43 bile acids were quantified using calibration curves and corrected by addition of internal standards. Bile acids that were not detected were assumed to be absent, and consequently set to 0 nmol/g. Only those BAs detected in at least 50% of the animals in at least one group were considered for downstream statistical analyses.

### Human samples

Participants who presented for a routine colonoscopy and/or underwent endoscopical polypectomy at the clinical department of the University Hospital Klinikum rechts der Isar (Technical University of Munich, Germany) were included in the ColoBAC registry (reference nr. 2018-322_8-S-SB) (29). For this work, individuals recruited between 2019 and 2022 were included: control, N = 8; hyperplastic polyp, N = 7; tubular adenoma, N = 9; carcinoma, N = 7. Lesion (abnormal) and adjacent healthy tissue (normal) was compared.

Proliferation in colorectal biopsies from patients who underwent screening colonoscopy at the University Hospital Dresden (Technical University Dresden, Germany) before and after treatment with Quantalan®, Colestyramin® or Cholestagel® between 2000 and 2024, was assessed retrospectively. Inflammation was evaluated by a board-certified pathologist at the Technical University Dresden. Studies were conducted with the approval and in compliance with the institutional guidelines and respective authorities at the Technische Universität Dresden (EK 249052019).

### Immunostaining

Human colon samples were routinely fixed in formalin, dehydrated, and embedded in paraffin. Representative areas of interest were annotated by a pathologist and tissue micro arrays (TMAs) with 1.0 mm core size and 1-3 replicates per patient were designed (**Supplementary Figure S2**). The TMAs includes 8 control patients (healthy mucosa), 18 tubular adenomas, 13 tubulo-villous adenomas, 4 hyperplastic polyps, 9 adenocarcinomas and paired, lesions-adjacent colonic mucosa. In addition, whole slides of 3 polyps, 3 adenocarcinomas, and adjacent normal mucosa were examined. Hematoxylin eosin staining was performed according to a standard protocol. Immunohistochemistry was performed either on a Ventana Benchmark XT platform (PDK4, FXR, Ki67) or on a Leica Bond RXm autostainer (TGR5, MUC2, GAL3ST2). Antibody binding was visualized chromogenically using 3,3’-diaminobenzidine. Details on the primary antibodies are: FXR (clone, OTI4F12; host, mouse; dilution, 1:75; supplier, Invitrogen, cat. no. MA5-27121); GAL3ST2 (polyclonal; host, rabbit; dilution, 1:50; supplier, Invitrogen, cat. no. PA5-64472); Ki67 (clone, Mib1; host, mouse; dilution, 1:50; supplier, Dako, cat. no. M724001); MUC2 (polyclonal; host, rabbit; dilution, 1:300; supplier, Novus Biologicals, cat. no. NBP1-31231); PDK4 (polyclonal; host, rabbit; dilution, 1:50; supplier, Invitrogen, cat. no. PA5-13776). Slides were then digitized by using a Leica AT2 slide scanning platform and image analysis, including semi-quantitative scoring was performed using Aperio ImageScope (Leica Biosystems Pathology Imaging, v12.4.6. For the markers MUC2, Ki67, and FXR, the frequency of cytoplasmic (MUC2) or nuclear (Ki67, FXR) positivity in epithelial cells was determined as a percentage (%). Cytoplasmic expression of TGR5, PDK4, and GAL3ST2 in epithelial cells was scored based on staining intensity grades (score 0-3).

## Results

To study the effects of secondary bile acid production on the gut epithelium under controlled conditions, we performed a gnotobiotic experiment. Germfree *Apc*^1638N/+^ mice were colonized with either the synthetic community OMM12 (20) alone or with addition of the species *E. muris* (OMM12+E), which produces secondary bile acids by 7α-dehydroxylation, mainly DCA (22). Epithelial cells isolated from the colon were analyzed by scRNA-Seq after 4 weeks of colonization.

### *E. muris* colonization triggered DCA production *in vivo*

Colonization profiles in the colon and bile acid concentrations in the cecum are shown in **Figure 1**. Nine of the 12 OMM strains colonized. *Bifidobacterium animalis*, *Limosilactobacillus reuteri*, and *Muribaculum intestinale* were not detected (**Figure 1a**). *Akkermansia muciniphila* and *Bacteroides caecimuris* were the two most dominant bacteria, whilst *Acutalibacter muris* and *Enterococcus faecalis* occurred at low relative abundances. *A. muris* did not colonize OMM12 mice, whereas the relative abundance of *E. faecalis* was higher in the OMM12+E group (5.5 *vs.* 0.15%), but results did not reach statistical significance. As expected, *E. muris* was detected only in the OMM12+E group with a mean relative abundance of 3.7 ± 1.2%. Bile acid concentrations in the cecum were characterized by marked differences between individual mice (**Figure 1b**). Only DCA was significantly different between the two colonization groups, as it was detected exclusively in the OMM12+E mice.

**Figure 1.**
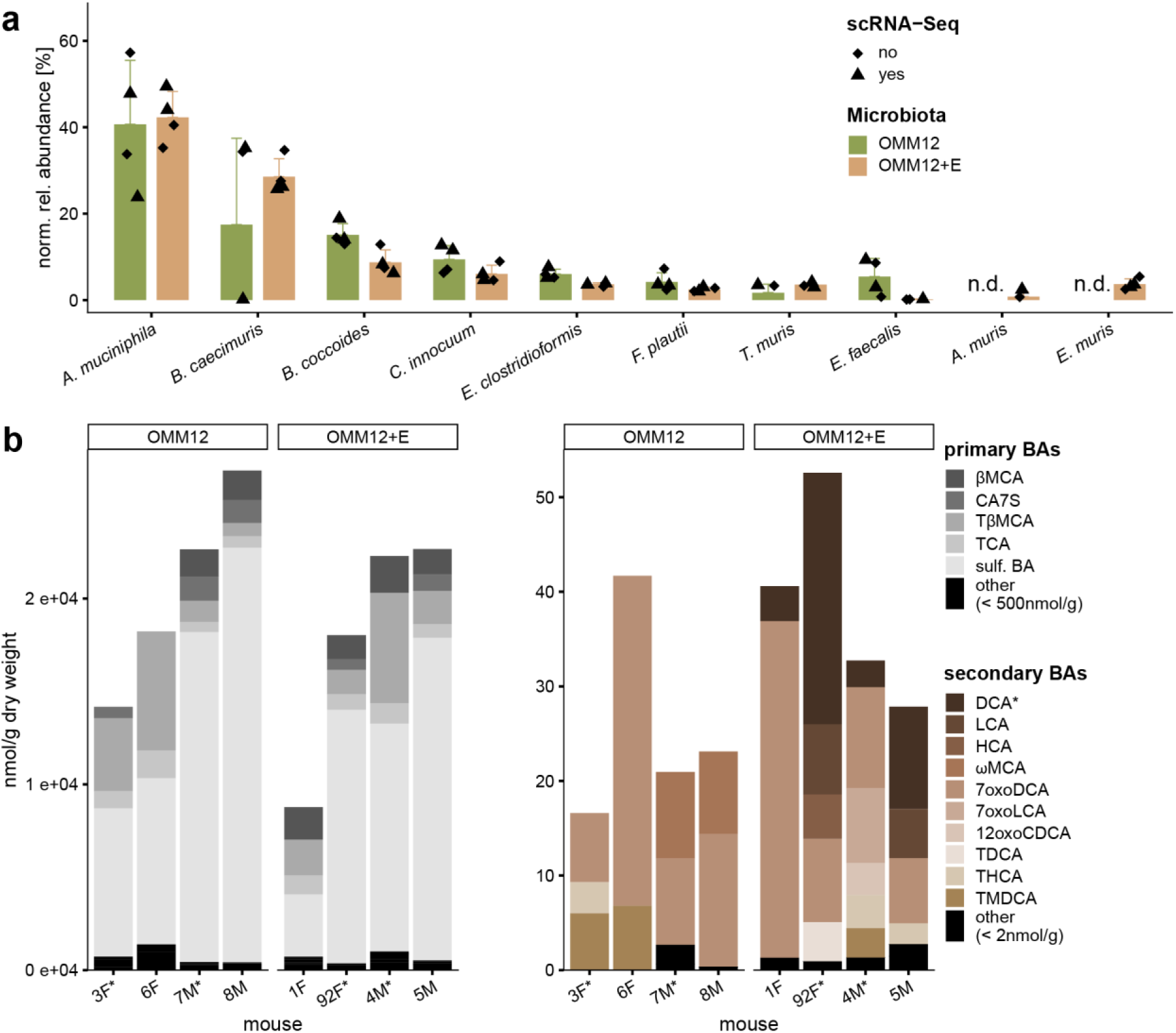
Microbiota and bile acid composition in the gut of gnotobiotic *Apc*^1638N/+^ mice. **a** Relative abundance of bacterial strains in the colon, as detected by 16S rRNA gene amplicon sequencing. OMM12, Oligo-Mouse Microbiota; OMM12+E, with addition of *E. muris* DSM 28560^T^. Triangles indicate the mice used for scRNA-Seq. **b** Composition of primary (left, grey) and secondary (right, brown) bile acids. The mice marked with a star below the x-axis were analyzed by scRNA-Seq.

### Epithelial cell landscape in the colon differed between the colonization groups

Next, we performed single cell RNA sequencing of epithelial cells isolated from the colon of OMM12 *vs.* OMM12+E mice. After processing and quality filtering of the scRNA-Seq data, 6,941 cells were analyzed in the OMM12 group, with an average of 8,914 sequenced molecules and 1,933 unique genes per cell. The OMM12+E sample contained a total of 5,112 cells, with 10,313 molecules and 1,993 unique genes per cell. A resolution of 0.3 was used for cell clustering, resulting in 12 distinct cell clusters (**Figure 2a**). When the distribution of cells was compared between the two colonization groups, fewer cells were observed within Cluster 5, 7, and 8 in the presence of *E. muris*, whilst Cluster 2 was populated with more cells in these mice (**Figure 2b**). Marker genes were used for cell type assignment, which resulted in the following classification of cell clusters (**Figure 2c**): Cluster 2, differentiated enterocytes located in the upper region of crypts (Caecam^high^, CD44-, Krt20+); Cluster 5, proliferating stem cells (Lgr5+, Mki67+); Cluster 7, goblet cells (Atoh1+, Spdef+) located in the middle region of crypts (Caecam^high^, CD44-); Cluster 8, almost absent in OMM12+E mice and characterized by increased proliferation (Mki67), is proposed to correspond to transit amplifying cells that develop into goblet cells of cluster 7.

**Figure 2.**
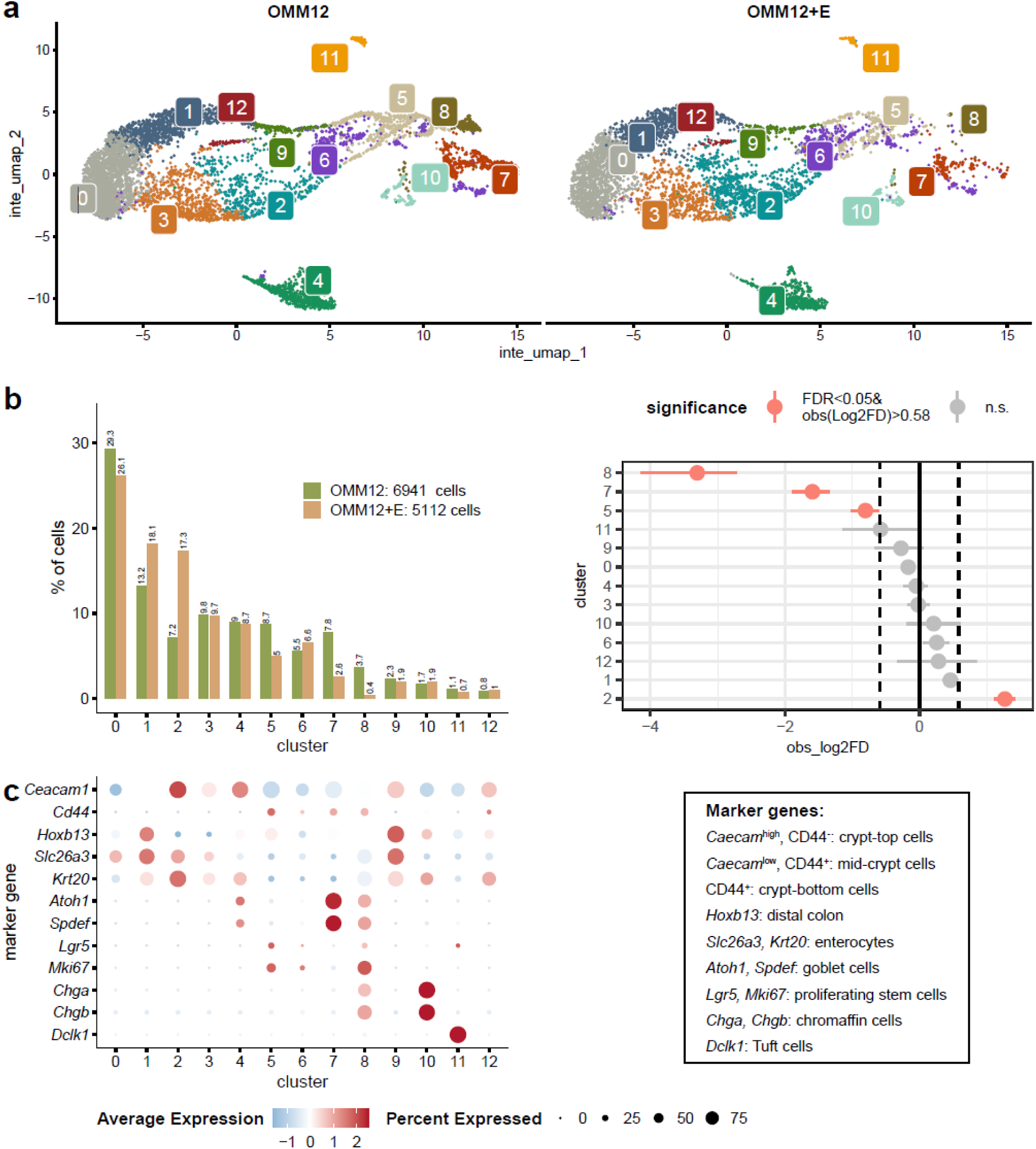
Transcriptomic analysis of colonic epithelial cells by scRNA-Seq. **a** UMAP of single cells by colonization group. Each plot (left: OMM12, Oligo-Mouse Microbiota 12; right: OMM12+E, with the addition of *E. muris*) represents a pool of colonic epithelial cells isolated from the colon of two mice. **b** Percentage of cells per cluster and corresponding over-/under-representation after permutation and bootstrapping. **c** Cell type identification based on marker genes (24).

### Differential gene expression in the colonic epithelium of *E. muris-*colonized mice

Since the main difference between the two colonization groups was secondary bile acid production, we examined the expression of bile acid-related genes. Absorptive enterocytes within cluster 2, with higher cell density in mice colonized with *E. muris* (see above), were characterized by high expression of the bile acid transporters ASBT (*Slc10a2*), MRP3 (*Abcc3*), OSTα (*Slc51a*), and OSTβ (*Slc51b*) (**Figure 3a**), and the nuclear bile acid-binding receptors PXR (*Nr1i2*), VDR (*Vdr*), and FXR (*Nr4h1*) (**Figure 3b**). The G-protein coupled receptor TGR5 (*Gpbar1*) was expressed mainly in chromaffin cells (cluster 10), which agrees with the literature (12).

**Figure 3.**
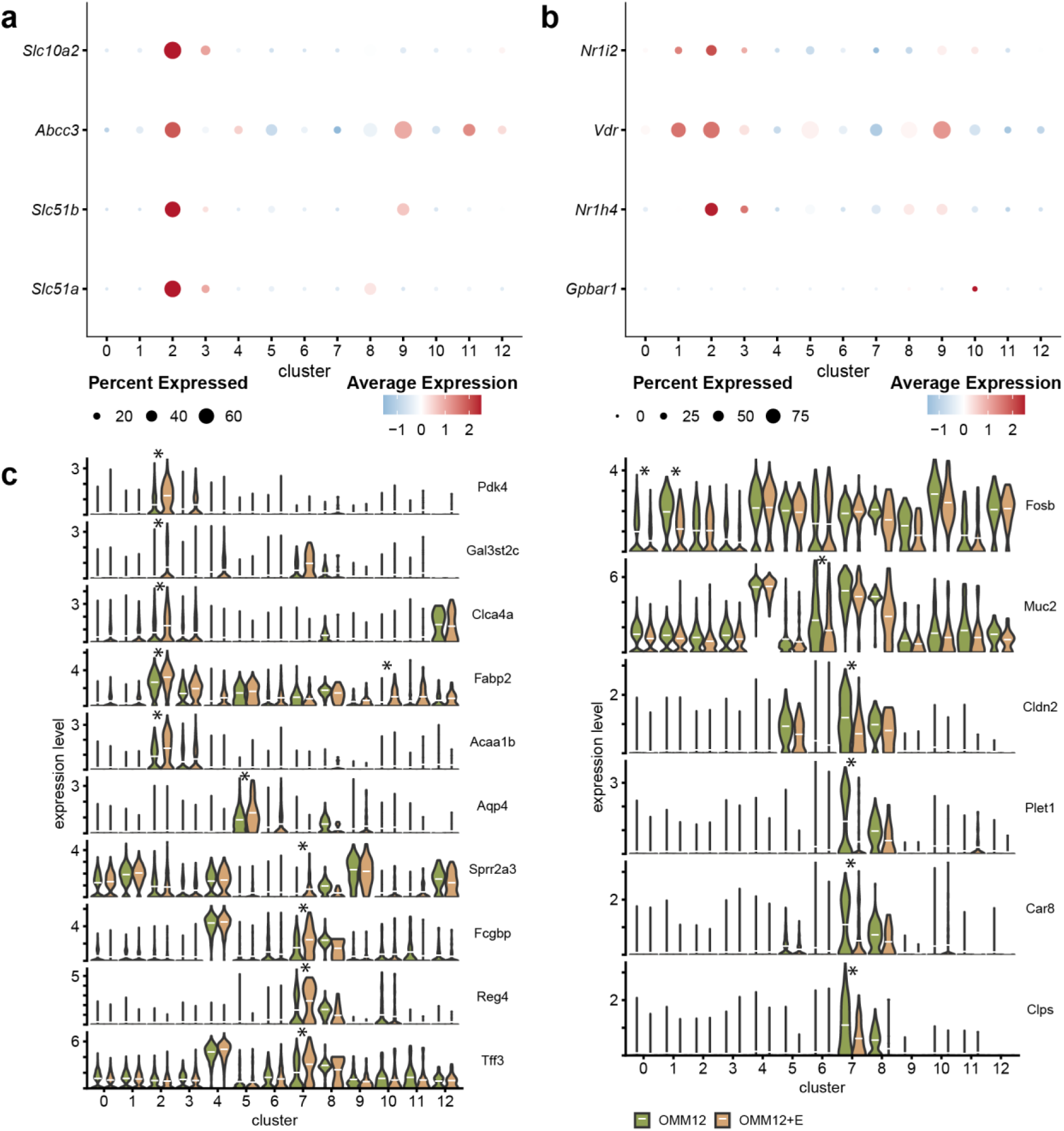
Bile acid-related and differentially expressed genes. **a** Genes encoding for the bile acid transporters ASBT (*Slc10a2*) on the apical side of cells, OSTα and OSTβ (*Slc51a* and *Slc51b*), and MRP3 (*Abcc3*) on the basolateral side. The dot plot was generated with scaled data (mean expression across the dataset is set to 0 with a standard deviation of 1). Negative average expression corresponds to expression below the mean expression of the dataset. **b** Genes encoding for bile acid receptors: PXR (*Nr1i2*), VDR (*Vdr*), FXR (*Nr1h4*), TGR5 (*Gpbar1*). In a and b, the average expression of the given gene (y-axis) between the different cell clusters (x-axis) is shown with a color gradient (red, high expression; blue, low expression). **c** Violin plots of differentially expressed genes as determined by the FindMarkers function of Seurat, based on Wilcoxon tests. Stars indicate highly (log-fold change >0.9) and significantly (adjusted p-value <0.05) affected genes with higher (left panel) or lower (right panel) expression in the OligoMM+E mice compared with the control mice without DCA-producing *E. muris*. The white horizontal lines show the mean gene expression per cluster/colonization. Gene designations: *Acaa1b*, acetyl-Coenzyme A acyltransferase 1B; *Aqp4*, aquaporin 4; *Car8*, carbonic anhydrase 8; *Clca4a*, chloride channel accessory 4A; *Cldn2*, claudin 2; *Fabp2*, intestinal fatty acid binding protein 2; *Clps*, colipase; *Fcgbp*, Fc fragment of IgG binding protein; *Fosb*, FosB proto-oncogene, *AP-1*, transcription factor subunit; *Gal3st2c*, galactose-3-O-sulfotransferase 2C; *Muc2*, mucin 2; *Pdk4*, Pyruvate dehydrogenase kinase 4; *Plet1*, placenta expressed transcript 1; *Reg4*, regenerating islet-derived family, member 4; *Sprr2a3*, small proline-rich protein 2A3; *Tff3*, intestinal trefoil factor 3.

Differential expression analysis within each cell cluster between the two mouse colonization groups revealed candidate genes with increased expression in the colon of OMM12+E vs. OMM12 mice, particularly in clusters 2 (bile acid-responsive enterocytes) and 7 (goblet cells) (**Figure 3c**). Pyruvate dehydrogenase kinase 4 (*Pdk4*), galactose-3-O-sulfotransferase 2C (*Gal3st2c*), chloride channel accessory 4A (*Clca4a*), intestinal fatty acid-binding protein 2 (*Fabp2*), and acetyl-Coenzyme A acyltransferase 1B (*Acaa1b*) were all significantly and highly (Log2FC >0.9) upregulated in cluster 2 cells. *Fabp2* was also significantly more expressed in cluster 10 (chromaffin cells) in the presence of *E. muris*. In addition, OMM12+E mice were characterized by higher expression of aquaporin 4 (*Aqp4*) in cluster 5 (stem cells) and of Fc fragment of IgG binding protein (*Fcgbp*), small proline-rich protein 2A3 (*Sprr2a3*), regenerating islet-derived family member 4 (*Reg4*), and intestinal trefoil factor 3 (*Tff3*) in cluster 7 (goblet cells). No significantly and highly (p.adj <0.05; Log2FC >0.9) upregulated genes were detected in the other cell clusters (0, 1, 3, 4, 6, 8, 9, 11, 12) due to *E. muris* colonization (**Supplementary Figure S3**).

In summary, the addition of *E. muris* to a synthetic community of mouse gut bacteria and the associated production of DCA *in vivo* was linked to changes in cell type composition and gene regulation in the colonic epithelium of *Apc*^1638N/+^ mice. Due to our previous findings on the causal role of DCA-producing bacteria, including *E. muris*, in promoting early-stage colorectal tumorigenesis (17), we used biopsies from patients with polyps or colorectal cancer to examine the expression of proteins corresponding to the top-regulated genes identified by scRNA-Seq.

### PDK4 and MUC2 expression was altered in abnormal human colonic tissue

Biopsies of normal mucosa or intestinal lesions (hyperplastic polyp, adenoma, carcinoma) from subjects undergoing endoscopy were analyzed by immunostaining to assess the expression of: (i) proteins encoded by candidate genes from the scRNA-Seq analysis or related to bile acid signaling: galactose-3-O-sulfotransferase 2C (GAL3ST2C), pyruvate dehydrogenase kinase 4 (PDK4), and mucin 2 (MUC2); and (ii) the bile acid receptors, farnesoid X-activated receptor (FXR) and G protein-coupled bile acid receptor 5 (TGR5) (**Figure 4**). Comparison of protein expression between tissues from control individuals or the patients (abnormal tissue) showed a decrease for FXR and MUC2 in the case of carcinoma (p < 0.01; Wilcoxon Rank Sum Test), indicating impaired bile acid signaling via FXR and abnormal mucus production (**Figure 4**). When assessing the differences between normal and abnormal tissue within a patient, PDK4 expression was increased in hyperplastic polyp and tubular adenoma, and MUC2 expression was decreased in tubular adenoma and carcinoma, mirroring the changes observed in colonic mucosa of the gnotobiotic *Apc*^1638N/+^ mice exposed to *E. muris* and DCA.

**Figure 4.**
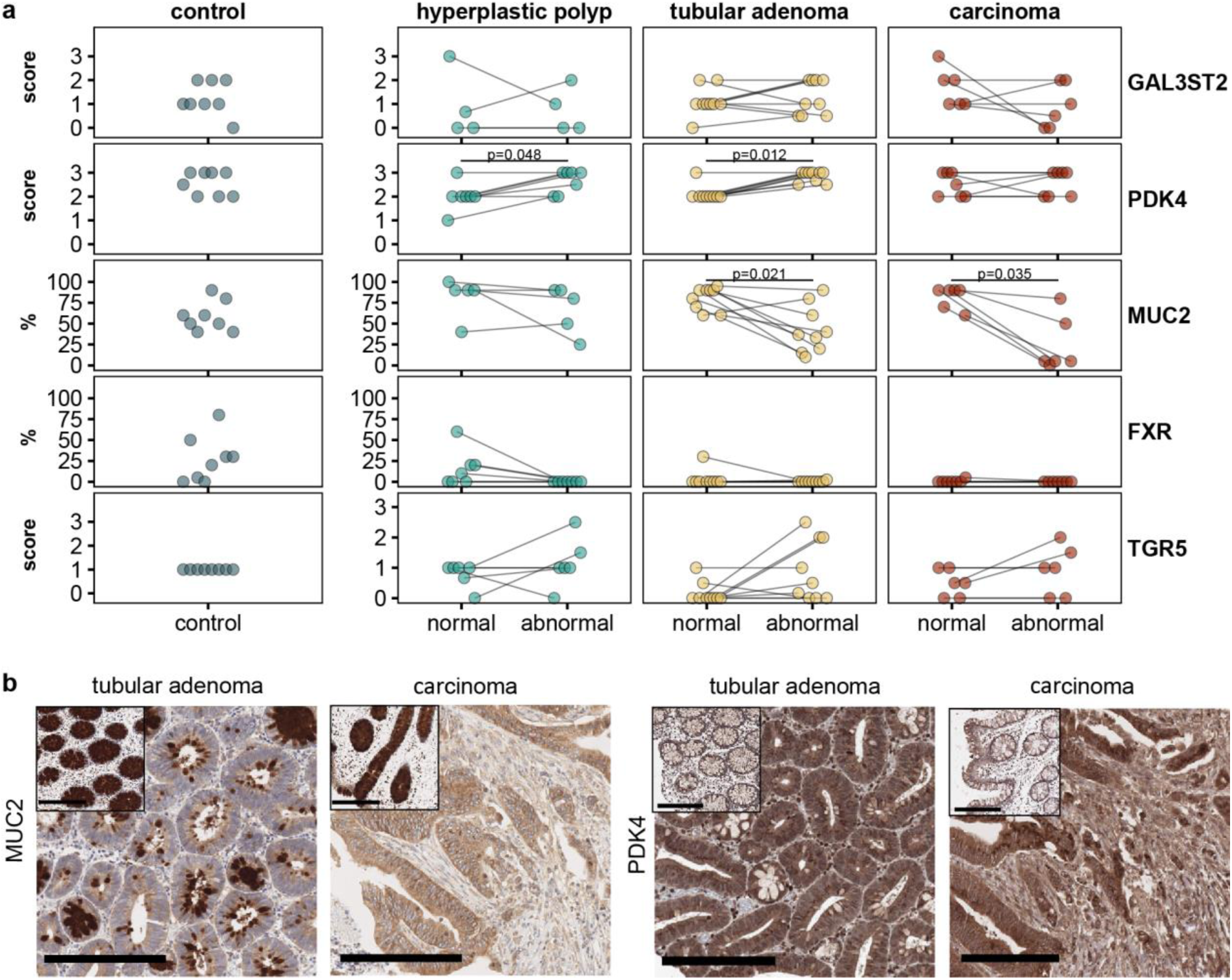
Candidate proteins stained in human colonic biopsies. All details about the patients, staining, and analysis are provided in the methods. The analysis included colonic tissue from individuals with hyperplastic polyp, tubular adenoma, or carcinoma. Tissue slides were assessed in a blinded manner by a pathologist and assigned a score (GAL3ST2, PDK4, TGR5) or fraction of cells positive for the given protein (MUC2, FXR). The dot plots show values in control subjects (left panel) and comparisons between paired biopsies in patients, *i.e.*, normal tissue adjacent to an abnormal region (right panel). Statistics: Paired Wilcoxon test with Benjamini-Hochberg correction. **b** Representative image of MUC2 and PDK4 staining. Cytoplasmic MUC2 staining of goblet cells was decreased in tubular adenoma (first image from left, 20 % positive) and rare in carcinoma (second image from left, 5 % positive) compared to healthy adjacent mucosa (90 % positive, insets) (scale bars = 200 µm). PDK4 showed a strong cytoplasmic signal in both, adenomatous and carcinomatous colonic epithelium (right images; score 3). In the adjacent normal colonic epithelium, PDK4 intensity was decreased (insets; score 2) (scale bars = 200 µm).

### Bile acid-scavenging drugs enhanced epithelial proliferation in the human colon

We previously found that DCA induces epithelial cell proliferation (Ki67 staining) and experimental tumorigenesis in the intestine of pigs and mice (17). Therefore, in addition to the experimental approach in gnotobiotic mice described above, which enabled the manipulation of DCA production to study epithelial responses under controlled conditions, we investigated the effects of clinically relevant modulation of intestinal bile acids on epithelial cell proliferation. Colon biopsies from patients treated with bile acid-scavenging drugs (colestyramine or cholestagel) due to ileal resection and chologenic diarrhea were examined by immunostaining for Ki67 before and after treatment. We observed a significantly higher proliferation of the epithelium in colonic tissue after treatment, regardless of the inflammatory status of the patient (**Figure 5**). A slight to marked increase in proliferation was observed in most of the patients (N = 12 of 17), while no change was observed in 4 patients and a slight decrease in 1 individual.

**Figure 5.**
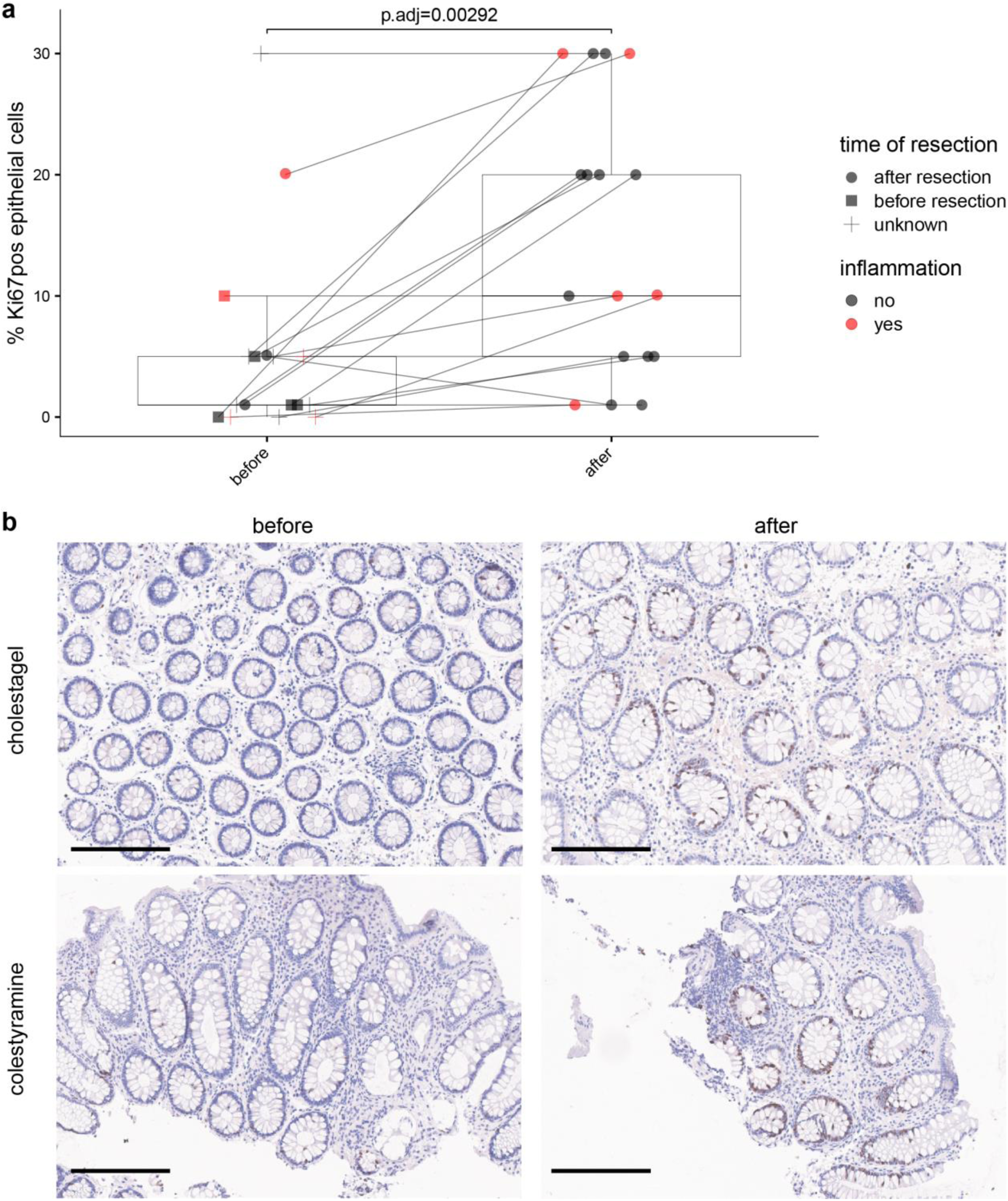
Effect of bile acid-scavenging drugs on epithelial cell proliferation in the human colon. This data included staining before and after an intervention with colestyramine or cholestagel. Ki67 was measured by immunostaining as described in detail in the methods. **a** Percentage of Ki67-positive (pos) cells before and after intervention. The black lines connect the two measurements for the same individual. Statistics: Paired Wilcoxon test with Benjamini-Hochberg correction. **b** Representative image of Ki67 staining. Scale bars, 200 µm.

## Discussion

Secondary bile acids produced by intestinal bacteria, especially DCA, play a causal role in the development of CRC. This is partly due to their effects on the gut epithelium, which need to be studied *in vivo*. Therefore, we performed gnotobiotic experiments with DCA-producing bacteria followed by scRNA-Seq and immunostaining of human colon tissue to assess changes in epithelial responses due to bile acid modulation.

Single-cell transcriptomics has emerged as a powerful method to dissect intestinal epithelial responses at high resolution (30). The effects of DCA on the gut epithelium have not yet been investigated using this technique. Previous work reported bulk-sequencing, transcriptomic data in whole colon tissue from female C57BL/6J mice treated daily with 400 µM DCA enemas for 7 days with or without dextran sulfate sodium treatment (31). DCA-induced changes were mainly related to cell growth and death, membrane transport, and lipid metabolism, as per KEGG analysis. The authors reported 808 up-regulated and 366 down-regulated genes, without providing further information. Our single-cell data from isolated colonic epithelial cells exposed to physiological concentrations of DCA in the gnotobiotic mice revealed that goblet cells and bile acid-sensitive absorptive enterocytes were the primary cell types affected by colonization. In enterocytes, *Pdk4* (pyruvate dehydrogenase kinase 4) (Gene ID: 27273), which encodes for a mitochondrial protein that inhibits the pyruvate dehydrogenase complex and thereby regulates glucose metabolism (32), was one of the most highly expressed genes compared with control mice. Gene expression of FABP2 (fatty acid binding protein 2) (Gene ID: 14079), which is involved in the transport and metabolism of long-chain fatty acids and in the modulation of cell growth and proliferation, was also increased in bile acid-sensitive enterocytes. These changes indicate alterations in cellular metabolism in the presence of DCA. Colon tumor tissue is known to have an altered lipidome compared with normal human mucosa (33). In addition, previous work reported increased *Pdk4* expression in gut tissue of tumor-prone mice (Mthfr^+/−^ BALB/c mice fed a folate-deficient diet) based on microarray analysis (34). The same authors found increased *Pdk4* expression in the normal mucosa of CRC patients (N=22) using qPCR. This corroborates our own findings: increased *Pdk4* expression in the colon of gnotobiotic mice exposed to DCA, which was mirrored by enhanced protein expression in abnormal colonic mucosa of patients at early to mid-stages of the disease (hyperplastic polyp and tubular adenoma). Other studies have reported the involvement of PDK4 in cellular migration and invasion, apoptosis, and epithelial cell extrusion (35, 36). In summary, PDK4 may be involved in the long-term tumorigenic effects of DCA produced by gut bacteria, but more work is needed to study the underlying molecular mechanisms.

We observed reduced cell density of goblet cells and transit amplifying cells in the colon of mice colonized with DCA-producing bacteria. In addition, *Muc2* expression was decreased in several cell clusters, reaching statistical significance in cells of cluster 6 (undefined). Previous *in vitro* work reported that DCA upregulated MUC2 expression in HM3 and HM3M2 cell lines (37). However, the DCA concentration (>100 µM) was higher than the physiological concentrations detected in our gnotobiotic mice (≤25 µM). We also observed that cluster 7 cells were characterized by decreased gene expression of claudin 2, indicating DCA-induced effects on cell-cell interactions (38). In contrast, *Reg4* expression was significantly increased in the presence of DCA. REG4 has been associated with aggressive CRC phenotypes and proposed as an early event in colorectal carcinogenesis (39). Furthermore, we observed that MUC2 expression was decreased in abnormal human colon tissue at a mid- to late stages of the disease (tubular adenoma and carcinoma). This is consistent with previous work in human tissues (40, 41). Considering that *Muc2*-deficient mice spontaneously develop rectal tumors (42), this data taken together may suggest that goblet cells are a main cell target of DCA, possibly contributing to CRC development when production is high due to consumption diets rich in fat over long periods of time.

DCA influences intestinal epithelial proliferation. However, data has been conflicting, even when considering only mechanistic studies and work performed *in vivo* or *ex vivo,* as discussed below. DCA inhibited mucosal healing in mice following biopsy-induced wounds in the colon, when administered at 3-4 mg/mouse by daily enemas for up to 6 days (14) and using an intestinal injury model in neonatal rats with a regimen of 5 mM three times daily by oral gavage for 4 days (43). In contrast, 5 µM DCA treatment *ex vivo* increased proliferation in biopsies of the normal sigmoid colon from six patients (with visible or occult fecal blood loss, abdominal discomfort, or diarrhea of unknown origin) as measured by bromodeoxyuridine immunohistochemistry (44). In addition, Sorrentino *et al.* (11) reported that bile acid-induced TGR5 activation increased intestinal epithelium proliferation and promoted intestinal stem cell renewal. In contrast to the overly negative effects attributed to DCA in the past, a new paradigm is that it regulates important physiological gut functions (*e.g.*, facilitating tissue regeneration) under steady-state conditions (*e.g.*, optimal diet), but becomes detrimental at higher concentrations under sub-optimal conditions (diet rich in fat), including increased epithelial cell proliferation which contributes to CRC development (17, 45). In this context, we sought to assess epithelial cell proliferation *in situ* after bile acid modulation, using colonic tissue from ileostomy patients treated with bile acid scavengers, which enhance bile acid excretion in stool. Strikingly, proliferation (Ki67 staining) was markedly increased after treatment, which may put this patient population at risk of developing CRC in the long term.

Taken together, this study provides new data on intestinal epithelial responses in relation to bile acid modulation and CRC. However, the work also has some pitfalls. Although our specific colonization strategy in gnotobiotic mice enabled the analysis of causal effects, we cannot exclude that the observed differences are not solely due to DCA, as the 7α-dehydroxylating species used, *E. muris*, may also exert additional functions in the mouse gut. In addition, we were not able to test all candidate genes from the scRNA-Seq data in human tissues due to either missing or non-functional antibodies. Finally, we were unable to determine bile acid concentrations in the patient samples due to the limited amount of tissue available. Therefore, we can only speculate that the increased proliferation observed in the colonic epithelium after treatment with bile acid scavengers is due to increased DCA production because of the large pool of primary bile acids entering the colon and available for bacterial transformation.

## Data availability

The 16S rRNA gene amplicon data from the gnotobiotic mice was submitted to the European Nucleotide Archive and is accessible via project accession no. PRJEB74082. The scRNA-Seq data were submitted to the NCBI and are accessible under BioSample accession SAMN40577731 and SAMN40577732 within BioProject PRJNA1090886.

## Acknowledgements

We are thankful to: (i) Susan A. Jennings (Functional Microbiome Research Group, Institute of Medical Microbiology, University Hospital of RWTH Aachen, Germany) for her outstanding support with gnotobiology; (ii) Aline Dupont and Stefan Schlößer (Institute of Medical Microbiology, University Hospital of RWTH Aachen, Germany) for their help with epithelial cell isolation; (iii) Johannes Schöneich (Institute of Medical Microbiology, University Hospital of RWTH Aachen, Germany) for helping with single-cell RNA sequence data analysis; (iv) the Flow Cytometry and Genomics Core Facilities of the Interdisciplinary Center for Clinical Research (IZKF) Aachen within the Faculty of Medicine at RWTH Aachen University, including funding by the German Research Foundation (DFG), project no. 439895892; (v) the iBio TUM tissue bank for excellent technical assistance.

## Funding

TC received funding from the German Research Foundation (DFG), project no. 453229399 (CL481/8-1) and project no. no. 403224013, SFB1382 Gut-liver axis. SZ, KS, MM, and TC received funding from the DFG, project no. 395357507, SFB1371 Microbiome Signatures. SZ and TC received funding from the German Ministry for Research and Education (BMBF): project Mi-EOCRC (KZ. 01KD2102D).

## Authors contributions

EW, TG, and CV performed experiments and analyzed data; EW, AS, SZ, KS, MM, and TC designed the studies; EW, TG, AS, KP, SZ, KS, MM, and TC interpreted data; EW and TC curated data; AS, KP, RBL, SZ, KS, MM, and TC provided access to essential resources and methods; SZ, KS, MM, and TC secured funding. EW created the figures. TC reviewed the figures. EW and TC wrote the manuscript. All authors have reviewed and approved the final version of the manuscript. TC coordinated the project.

## Conflict of interest

The authors declare no conflicts of interest

---

**Supplementary Figure S1** Descriptive scRNA-Seq data on quality

**Supplementary Figure S2** Experimental scheme of the work with tissue microarrays (TMAs). Abbreviations: ID, identity; FFPE, formalin-fixed paraffin-embedded.

**Supplementary Figure S3** Volcano plot of differently expressed genes in the gut epithelium of mice colonized with the Oligo-Mouse Microbiota 12 (O) without or with addition of the deoxycholic acid (DCA)-producing species *Extibacter muris* (E). The data is shown per cell cluster.

## Notes

### Competing Interest Statement

The authors have declared no competing interest.

