## Supplementary figures and images for "Modulation of intestinal bile acids influences colonic mucosal responses"

### Suppl Fig S1

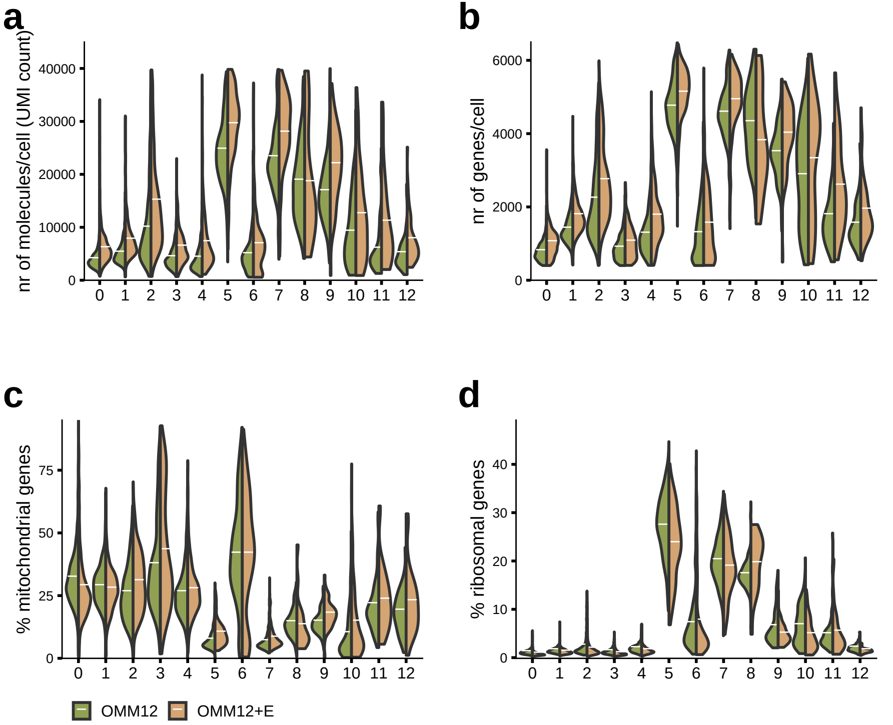

### Suppl Fig S2

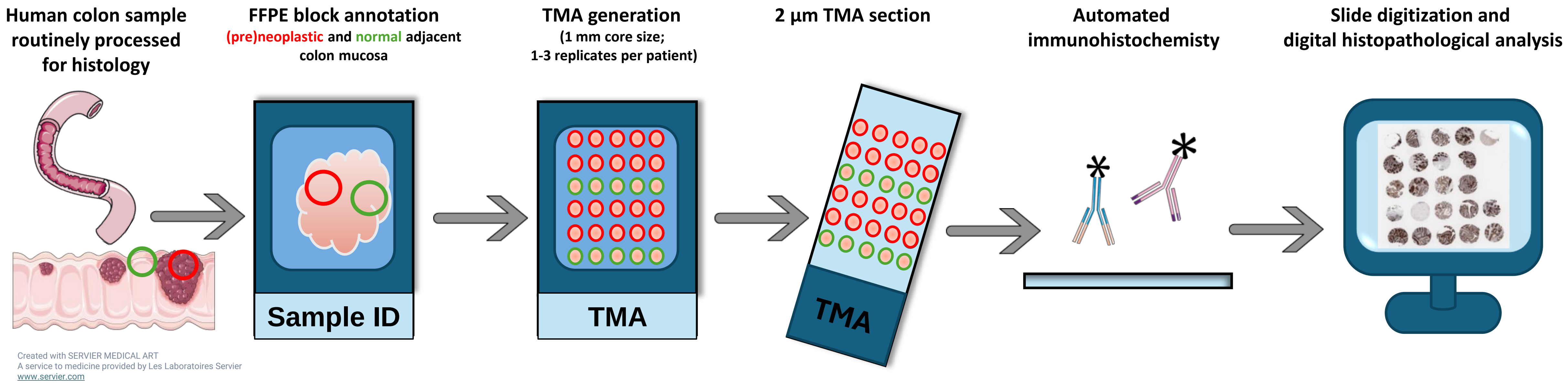

### Suppl Fig S3

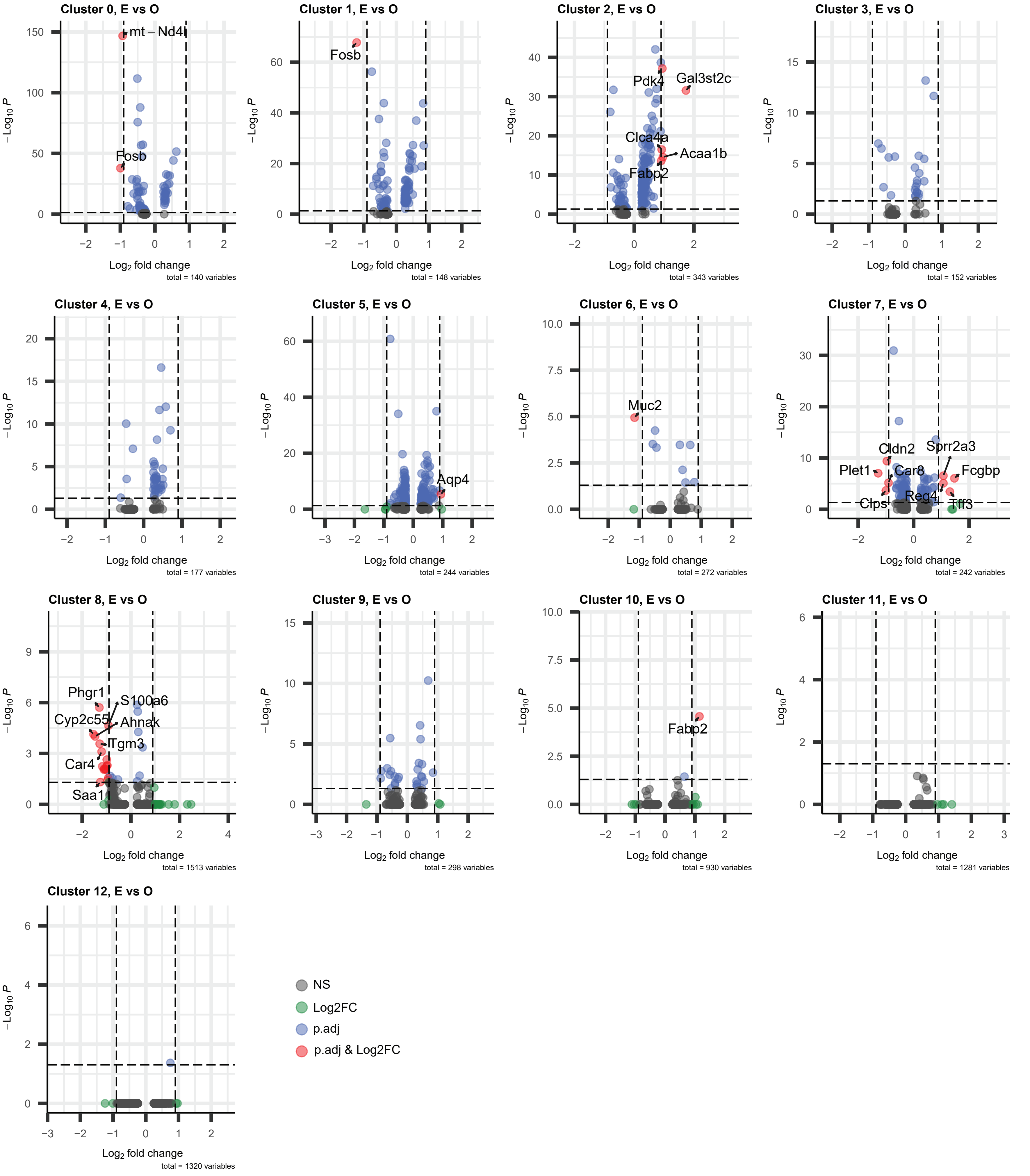
